# Geometricus Represents Protein Structures as Shape-mers Derived from Moment Invariants

**DOI:** 10.1101/2020.09.07.285569

**Authors:** Janani Durairaj, Mehmet Akdel, Dick de Ridder, Aalt DJ van Dijk

**Affiliations:** Bioinformatics Group, Department of Plant Sciences, Wageningen University and Research; Mathematical and Statistical Methods - Biometris, Department of Plant Sciences, Wageningen University and Research

## Abstract

**Motivation:** As the number of experimentally solved protein structures rises, it becomes increasingly appealing to use structural information for predictive tasks involving proteins. Due to the large variation in protein sizes, folds, and topologies, an attractive approach is to embed protein structures into fixed-length vectors, which can be used in machine learning algorithms aimed at predicting and understanding functional and physical properties. Many existing embedding approaches are alignment-based, which is both time-consuming and ineffective for distantly related proteins. On the other hand, library- or model-based approaches depend on a small library of fragments or require the use of a trained model, both of which may not generalize well.

**Results:** We present Geometricus, a novel and universally applicable approach to embedding proteins in a fixed-dimensional space. The approach is fast, accurate, and interpretable. Geometricus uses a set of 3D moment invariants to discretize fragments of protein structures into shape-mers, which are then counted to describe the full structure as a vector of counts. We demonstrate the applicability of this approach in various tasks, ranging from fast structure similarity search, unsupervised clustering, and structure classification across proteins from different superfamilies as well as within the same family.

**Availability:** Python code available at https://git.wur.nl/durai001/geometricus

## 1 Introduction

The number of structures added to the Protein Data Bank (Bernstein *et al*., 1977) has been increasing rapidly, with over 10,000 structures deposited in 2019 alone. Meanwhile, major advances have been made in the areas of homology-based and *de novo* protein structure modelling (Senior *et al*., 2020). This increased availability of protein structures has enabled protein biologists and bioinformaticians to start including structural data and information in protein function studies instead of being confined to the sole use of sequence data. These studies address a variety of questions, such as finding remote protein homologs with a similar structural fold, or defining the properties of a single protein family. Protein structures evolve slower than sequences, and encode long-range contact and fold information that are often crucial for protein activity. Hence, our understanding of molecular biology can be greatly enhanced by the inclusion of protein structures.

For both structures and sequences, choosing the right computational method to generate a representation of a protein for comparison and prediction purposes is crucial. This is especially true for machine learning methods, which often require variable-length sequences of amino acids, coordinate sets or other residue descriptors to be transformed into fixed-length representations. These representations can be used as input for supervised and unsupervised machine learning methods or be compared using standard vector distance formulae. As proteins typically cover a wide range of shapes, sizes and topological folds, the choice of representation is not always straightforward and may depend on the scale of the study. For instance, research questions addressing proteins within a single family may opt to use alignment-based representations (Simossis *et al*., 2003; Ma and Wang, 2014). These have the advantage of easy interpretability, as each residue can be directly mapped to a column in the transformed representation. However, alignment is computationally expensive and its accuracy decreases with decreasing protein similarity.

To solve this, alignment-free methods were introduced, which learn a reduced and condensed representation of proteins without an explicit alignment. There are many examples of such approaches using machine learning and deep learning methods to learn generic patterns and features of the protein sequence space (Alley *et al*., 2019; Rao *et al*., 2019). Structure-based representations also exist (Budowski-Tal *et al*., 2010; Liu *et al*., 2018b) but are generally more difficult to generate due to the three-dimensional nature of structures compared to the one-dimensional sequences. Some structure-based “alignment-free” methods generate a representation of a protein of the same length as the sequence and then use sequence alignment or calculate sequence similarity to compare these structural sequences in 1D (Lo *et al*., 2007; Le *et al*., 2009). The conserved nature of protein structure circumvents the problem of decreasing accuracy of sequence alignment in these approaches.

Many structure embedding techniques make use of a library of small structural fragments to which fragments of each input structure are compared, usually requiring the calculation of rotations and translations that would orient the input fragment and the library fragment in the same position (Budowski-Tal *et al*., 2010). To reduce the computational load of these structure-structure comparisons, library sizes are limited. Newer techniques which make use of deep learning (Liu *et al*., 2018b), do not need a library but still require a pre-trained model to generate new embeddings. In both cases the size, scale and resolution of the embedding is highly dependent on the initial choice of library fragments or training data used, and thus may not be applicable to research questions about a different protein set. Also, in both these approaches it is difficult to link predictive importance to functionally important residues or structural regions, which is often desired in studies aimed at understanding underlying mechanisms in protein biology.

To address these disadvantages of existing approaches, we introduce Geometricus, a novel structure embedding technique based on 3D rotation-invariant moments. Moment invariants were proposed in the 1960s for 2D images (Hu, 1962) and have been used extensively in the image processing field for object detection (Rizon *et al*., 2006) and character recognition (Flusser and Suk, 1994) among other applications. In the 1980s they were subsequently adapted to the 3D field (Sadjadi and Hall, 1980), and found applications in the fields of robotics (Se *et al*., 2001), gesture recognition (Kratz and Rohs, 2011), brain morphology (Mangin *et al*., 2004), and even in structural biology (Sommer *et al*., 2007). Rotation-invariant moments yield identical output when performed on any translated and/or rotated version of a set of continuous or discrete three-dimensional coordinates. This implies that coordinate sets can be compared without the need for superposition. Alternatively, any set of coordinates can be represented by a number of these moments.

To generate a Geometricus embedding for a protein structure, we fragment the structure into overlapping *k*-mers based on sequence, as well as into overlapping spheres, calculated for a certain radius, based on 3D coordinate information. Moment invariants are calculated for each of the coordinate sets corresponding to these structural fragments, and then binned into *shape-mers*, each of which represent a set of similar structural fragments. Counting the occurrences of these shape-mers across the protein yields a representation of the whole protein structure as a fixed-length vector of counts, similar to an amino acid *k*-mer count vector describing a protein sequence. As the moment invariant calculation is simple, the entire embedding process runs in the order of tens of milliseconds per protein and is easily parallelized. In addition, each element in the count vector can be mapped back to the residues forming the corresponding shape-mer, allowing for interpretation of predictive residues on par with alignment-based approaches. The shape-mer binning process is easily controllable, allowing for coarse shape-mer definitions for divergent proteins with distinct structures, or a fine-grained resolution for closely related proteins from the same family. This makes Geometricus suitable for a variety of tasks where library-based or model-based embeddings would struggle or require expensive retraining.

We demonstrate the effectiveness and versatility of Geometricus embeddings in a variety of machine learning approaches and other applications applied to datasets of varying structure similarity. Geometricus can be used for very fast structure similarity searches, while maintaining accuracy close to that obtained by alignment-based methods. The innate simplicity of the approach enables flexibility in application, such that embeddings can be optimized for the task at hand, as we demonstrate using datasets with proteins from different superfamilies and within the same family. Geometricus is available as a Python library at https://git.wur.nl/durai001/geometricus.

## 2 Methods

### 2.1 Protein Embedding

To generate embeddings for a set of proteins, we define so-called shape-mers which are analogous to sequence *k*-mers. A shape-mer represents a set of similar structural fragments, each a collection of coordinates in 3D space. The following sections describe the process of generating these structural fragments, their subsequent conversion into rotation- and translation-invariant moments, the moment-based grouping of structural fragments into shape-mers, and finally, shape-mer counting to obtain the resulting embedding.

#### 2.1.1 Protein Fragmentation

We consider two different ways of dividing a protein with *l* residues into structural fragments, using its *α*-carbon coordinates, 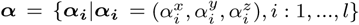.

1. ***k*-mer based** - for a given value of *k*, a protein is divided into *l k*-mer-based structural fragments, 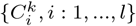 where

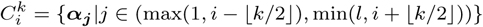 Here ⌊⌋ converts a floating point number to the closest integer value below it.
2. **radius based** - for a given radius *r*, a protein is divided into *l* radius-based structural fragments 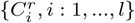 where

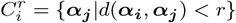

with *d*(***α***_***i***_, ***α*** _***j***_) being the Euclidean distance between ***α***_***i***_ and ***α***_***j***_. Practically, this is accomplished by constructing a KD-Tree on ***α***, using the KD-tree implementation in ProDy v1.10.11(Bakan *et al*., 2011) and querying by radius with each ***α***_***i***_ as the center.

While the *k*-mer based approach is effective in describing structural fragments that are sequential in nature, such as *α*-helices and loops, the radius-based approach can capture long-range structural contacts as seen in *β*-sheets, as well as distinct interaction patterns in space, as found in enzyme active sites. Both fragmentation methods have *𝒪* (*l*) time complexity.

Each resulting structural fragment is then transformed into four moment invariants, described in the next section. In our examples and results section we use a *k* of 16 and a radius *r* of 10 Å as a compromise between specificity of the structural fragments and effectiveness of the moment invariants. In principle, optimization of these parameters could lead to further improvements of our approach for specific applications, but we leave this open for future exploration.

#### 2.1.2 Moment Invariants

Three-dimensional moment invariants are computed using the formula of the central moment, defined below for a discrete set of *c* coordinates, with 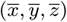 being the centroid:

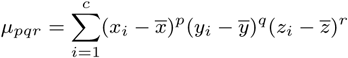

Using this formula, we then compute four moments that were previously used in a structural bioinformatics study to describe enzyme active sites (Sommer *et al*., 2007). These include the three second-order rotation invariants (*O*_3_, *O*_4_, and *O*_5_) described by Mamistvalov (1998) and a fourth invariant, *F*, described by Flusser *et al*. (2003). These four moment invariants are defined below:

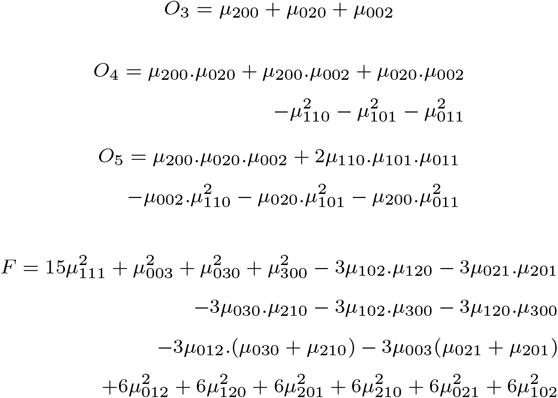

Thus, any structural fragment can be represented by a vector (*O*_3_, *O*_4_, *O*_5_, *F*). Moment invariant calculation is implemented using Numba v0.48.0 (Lam *et al*., 2015) and has 𝒪 (*c*) time complexity which is negligible for small values of *c*, as seen for *k*=16 (i.e. a maximum *c* of 16) and *r*=10 (*c* = 18 *±* 6).

#### 2.1.3 Discretization to Shape-mers

While the moment invariants obtained for each structural fragment can be directly compared, discretizing them enables collecting sets of fragments that resemble each other across multiple proteins. We convert the continuous and real-valued moment invariants to discrete shape-mers as follows:

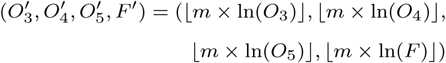

Here *m* is the resolution parameter, which defines the coarseness of the shape-mers, with higher values leading to more fine-grained separation of structural fragments. Thus, a shape-mer is defined by four discrete numbers and can describe any number of structural fragments. Figure 1 gives examples of moment invariant and shape-mer calculations (with *m* = 1) for three synthetic coordinate sets generated with the equation {***α***_***i***_ = (*R* cos(*i*), *R* sin(*i*), *i*), *i* : 1, *…*, 16} for *R* = 0, 0.5, and 2 respectively, each rotated by *±*45°, and translated by *±*10Å along the *x*-axis.

**Fig. 1.**
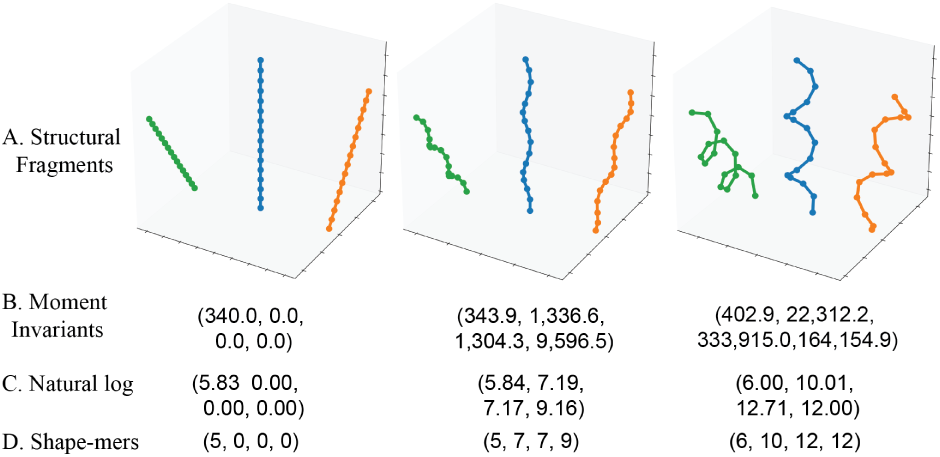
A. Three synthetic structural fragments, with a rotation of *±*45° and translation of *±*10Å between the middle fragment and the two outer ones. B. Moment invariants (*O*_3_, *O*_4_, *O*_5_, *F*) for each fragment. The three rotated and translated versions have the same moment invariant values. C. The natural log-transformed versions of (*O*_3_, *O*_4_, *O*_5_, *F*) and D. shape-mers 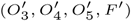 for each fragment.

#### 2.1.4 Counting Shapes

Given a set of *n* proteins, we generate a collection of shape-mers for each protein. The total number of shape-mers *s* is then the number of distinct shape-mers observed across all *n* proteins. A count vector of length *s* is calculated for each protein with each element recording the number of times the corresponding shape-mer appears in that protein. This counting is done separately for the *k*-mer and radius based approaches, as they represent different types of structural fragments. The two resulting count vectors are concatenated to form the final protein embedding. The entire embedding process has a time complexity of *𝒪* (*nl*) and takes around 50 milliseconds CPU time for proteins of medium length (400-600 residues). Note that different values for *m* (the resolution parameter) and different input sets of proteins will lead to different sets of shape-mers and embedding sizes. This allows the user to generate feature spaces tailored to the problem at hand.

### 2.2 Datasets

We apply Geometricus to a number of datasets to demonstrate the wide applicability of shape-mer based protein embedding. These are described below. The remaining sections use the acronyms defined here to refer to these datasets.

1. **CASP11** - 87,573 protein structures from the Critical Assessment of protein Structure Prediction XI (Moult *et al*., 2016) training set, obtained from the ProteinNet data source (AlQuraishi, 2019).
2. **CATH20** - The CATH database of protein structures (Pearl *et al*., 2003) categorizes proteins hierarchically based on secondary structure class (C), architecture (A), topology (T), and homology (H). From the CATH hierarchy, we selected 3,673 proteins with *<*20% sequence identity to each other from the top five most populated CAT categories. Table 1 shows the number of proteins per CAT category.
3. **SCOP-Lo** - The Structural Classification of Proteins (SCOP) database (Murzin *et al*., 1995) provides a detailed classification of structures based on their topologies and folds. We adapted the SCOP-Lo dataset from Lo *et al*. (2007). This dataset comprises 23,912 target proteins from ASTRAL SCOP 1.67 further divided into sets with 10%, 30%, 70% and 100% maximum sequence identity within each group respectively. It also contains a query set of 83 proteins each with at least two proteins from the same SCOP family in the 10% target protein set, and <10% sequence identity to other proteins in the query set.
4. **Pfam10** - The Protein families database (Pfam) (Bateman *et al*., 2002) collates a large set of protein families. Out of the twenty most populated Pfam domains, the ten accessions with most available structures are considered, resulting in a total of 3,053 structures. Table 2 lists the number of proteins for each of these ten Pfam accessions.
5. **CMGC** - 1,822 human protein structures in the CMGC kinase family were collected from the Kinase-Ligand Interaction Fingerprints and Structures (KLIFS) database (Kooistra *et al*., 2016). These are further divided into 660 cyclin-dependent kinases (CDK), 527 mitogen-activated protein kinases (MAPK), 268 casein kinase 2 (CK2) proteins, 160 dual specificity Tyrosine regulated kinases (DYRK), 122 glycogen synthase kinases (GSK), 61 cdc2-like kinases (CLK), 16 serine/threonine-protein kinases (SRPK), and 8 cyclin-dependent kinase-like kinases (CDKL).
6. **MAPK** - The 527 MAP kinases from the CMGC dataset are considered separately. These comprise 271 p38 MAPK structures (p38), 147 extracellular signal-regulated kinases (ERKs), and 109 c-Jun N-terminal kinases (JNKs).

Note that the low sequence identity between the proteins in many of these datasets clearly underlines the need for structure-based embedding.

**Table 1.** Number of proteins in each CAT category in the CATH20 dataset

| Class | Architecture | Topology | No. of Proteins |
| --- | --- | --- | --- |
| Mainly Alpha | Orthogonal Bundle | Arc Repressor Mutant, subunit A | 465 |
| Mainly Beta | Sandwich | Immunoglobulin-like | 700 |
| Mainly Beta | Sandwich | Jelly Rolls | 401 |
| Alpha Beta | 3-Layer (aba) Sandwich | Rossmann Fold | 1,660 |
| Alpha Beta | 2-Layer Sandwich | Alpha-Beta Plaits | 447 |

**Table 2.** Number of proteins for each Pfam accession in the Pfam10 dataset

| Pfam Accession | Short name | Description | No. of Proteins |
| --- | --- | --- | --- |
| PF00005 | ABC_tran | ABC transporter | 187 |
| PF00069 | Pkinase | Protein kinase domain | 438 |
| PF00076 | RRM_1 | RNA recognition motif. (a.k.a. RRM, RBD, or RNP domain) | 269 |
| PF00096 | zf-C2H2 | Zinc finger, C2H2 type | 144 |
| PF00400 | WD40 | WD domain, G-beta repeat | 906 |
| PF00440 | TetR_N | Bacterial regulatory proteins, tetR family | 164 |
| PF02518 | HATPase_c | Histidine kinase-, DNA gyrase B-, and HSP90-like ATPase | 87 |
| PF12796 | Ank_2 | Ankyrin repeats (3 copies) | 263 |
| PF13561 | adh_short_C2 | Enoyl-(Acyl carrier protein) reductase | 273 |
| PF13855 | LRR_8 | Leucine rich repeat | 322 |

### 2.3 Visualization of shape-mers and Geometricus embeddings

To visualize two commonly occurring *k*-mer-based shape-mers from the CASP11 dataset, we first randomly selected 1,000 structural fragments described by them. From these 1,000, one fragment was randomly chosen as the base and the remaining were superposed to the base using the Kabsch algorithm (Kabsch, 1976). For each fragment, the best superposition to the base of that fragment and its flipped version, in terms of the minimum Root Mean Square Deviation (RMSD), is taken in order to account for 360*°* rotations. We also visualized two radius-based shape-mers using two of their structural fragments shown in the context of their respective protein structures.

The Geometricus embeddings of the Pfam10, CMGC, and MAPK datasets, generated for different values of the *m* parameter, were reduced to two dimensions using the Python implementation (v0.3.10) of the Uniform Manifold Approximation and Projection (UMAP) algorithm by McInnes *et al*. (2018), with the cosine similarity metric and default settings.

### 2.4 Structure similarity search

We demonstrated how Geometricus can be used in structure-based similarity searches by applying it to the CATH20 and SCOP-Lo datasets. A pair of proteins is called similar if they share the same CAT category for the CATH20 dataset or the same SCOP category for the SCOP-Lo dataset, and dissimilar otherwise.

Typically, in structure similarity search applications, similarity scores are calculated for a small set of *query* proteins against a larger predefined and preprocessed *target set* of structures. Here, the target set determines which collection of shape-mers will be used in the search. For the CATH20 dataset, 70% of the proteins are randomly chosen as the target set. The SCOP-Lo dataset already has four defined target sets (10%, 30%, 70%, and 100% sequence redundant sets) which are each evaluated separately.

The pairwise similarity measure between two proteins is defined as the cosine similarity of their Geometricus embedding vectors, constructed with a low resolution (*m* = 0.25) to reflect the major structural differences expected between proteins in these two datasets. Proteins with a similarity score above a threshold *t* are predicted to be similar and those below *t* are predicted as dissimilar. We calculated similarity scores for all CATH20 proteins against the CATH20 target set, and the 83 SCOP-Lo query proteins against each of the four sequence redundant SCOP-Lo target sets. ROC-AUC curves were constructed by varying *t* to evaluate the correctness of the similarity search in these five cases.

### 2.5 CATH classification

A *k*-nearest neighbor classifier from the scikit-learn python library v0.22.1 (*k*=5, metric=“cosine”) was trained to predict the CAT category for the proteins in the CATH20 dataset with 50% of the data randomly chosen for training and the remaining for testing. We repeated this five times and report the average accuracy.

### 2.6 MAP kinase classification

To demonstrate the applicability of Geometricus for interpretable machine learning on protein structures, we performed classification on the MAPK dataset to predict the type of MAP kinase (namely p38, ERK, or JNK) from protein structure. This was accomplished using the decision tree classifier from the scikit-learn Python library (v0.22.1) (Pedregosa *et al*., 2011), with a random 70%-30% split of training and test data. The top two most predictive shape-mers from the trained classifier were then mapped back to the residues that they correspond to and visualized on one p38 structure (PDB ID: 3QUE), one ERK structure (PDB ID: 2OJJ) and one JNK structure (PDB ID 4KKG) using PyMOL (DeLano *et al*., 2002).

## 3 Results

### 3.1 Shape-mers capture common structural fragments across protein structures

We performed moment-invariant and shape-mer calculations on over 87,000 proteins in the CASP11 dataset to understand their distributions and patterns found across structurally divergent proteins. Figure 2A shows the log-distribution of each of the four moment invariants for the *k*-mer- and radius based structural fragmentation approaches. The radius-based approach shows wider distributions in general, which can be expected: different locations in a protein have different densities of residues leading to differing numbers of coordinates in the radius-based approach, while the *k*-mer based approach largely produces fragments with *k* coordinates except for some shorter fragments at the N- and C-terminal ends of each protein.

**Fig. 2.**
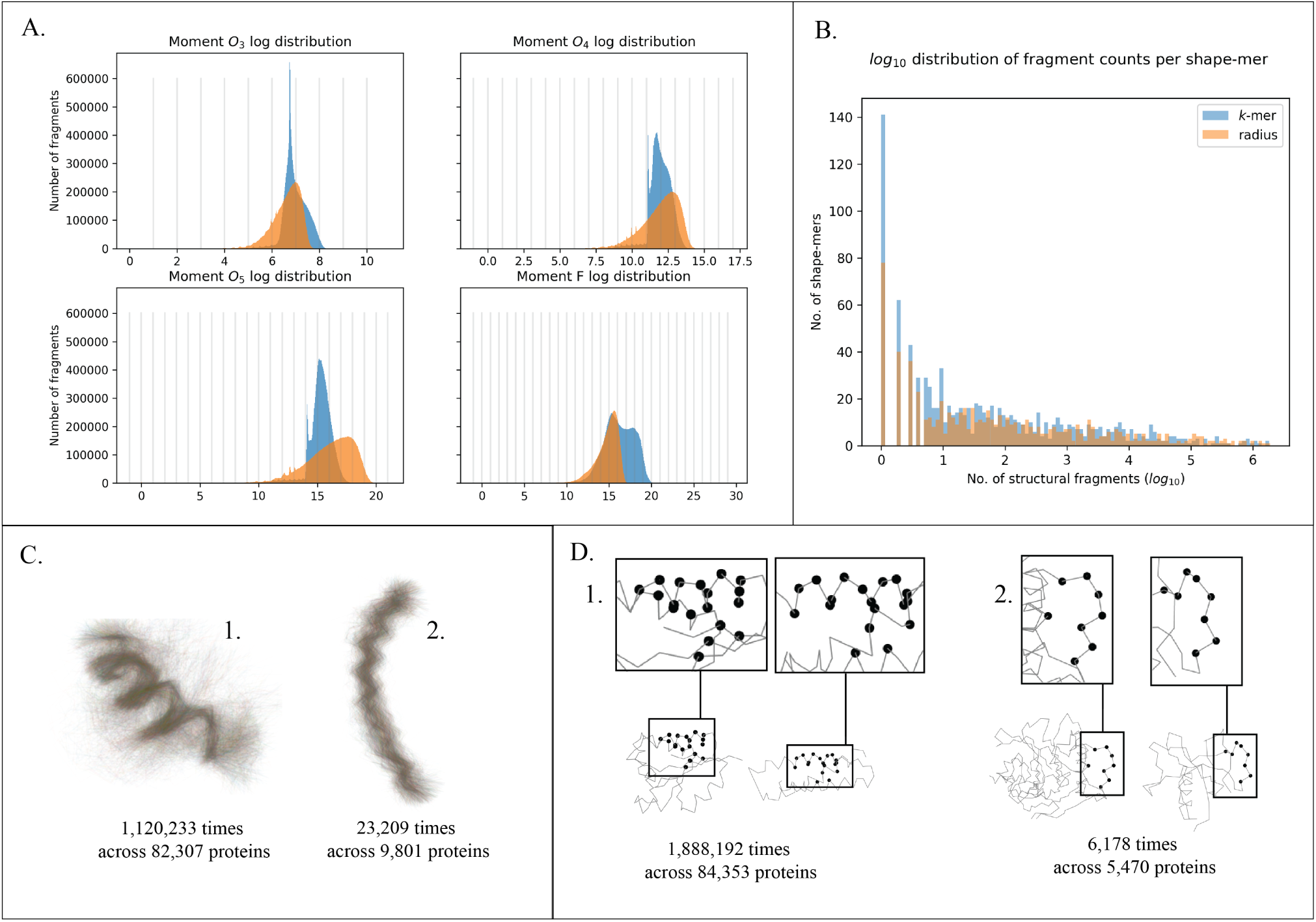
A. Natural log distribution of each of the four moment invariants across *k*-mer based structural fragments (in blue) and radius-based structural fragments (in orange) from proteins in the CASP11 dataset. B. Histogram of (log_10_ transformed) counts of fragments described by each shape-mer. C. Two examples of *k*-mer based shape-mers, shown using a thousand randomly selected fragments, superposed. D. Two examples of radius-based shape-mers, magnified, and highlighted as black dots on two protein structures in which they are found.

Shape-mers were computed from the moment invariants using a resolution *m* of 1 (see Methods). The resulting 565 *k*-mer shape-mers and 703 radius shape-mers do not all represent the same number of structural fragments. Figure 2B shows the log_10_ distribution of structural fragment counts represented by each shape-mer. Some shape-mers, at the right end of Figure 2B, are found over a million times, and unsurprisingly represent common structural fragments such as short, well-defined *α*-helices. One *k*-mer based and one radius-based example are shown in Figure 2C1 and Figure 2D1 respectively, both found across most of the proteins in the CASP11 dataset. Conversely, the shape-mers on the very left end of the Figure 2B represent only one structural fragment, likely loops or specific folds which are structurally and functionally unique and thus rare. The remaining shape-mers describe anywhere between one and a million fragments and may be specific to certain superfamilies or families of proteins. of Figure 2C2 shows an extended roll-like shape-mer found in almost 10,000 proteins and Figure 2D2 shows a sparse radius shape-mer found on the surfaces and ends of 5,000 proteins.

### 3.2 Geometricus can be used for fast and accurate structure similarity search and topology classification

A common application of structure-based embeddings is to perform a fast similarity search for an input structure across a database of structures and return the most similar candidates. We demonstrate the performance of Geometricus on this task using CATH and SCOP classifications as a ground truth measure of protein similarity.

Figure 3 shows the Receiver Operating Characteristic (ROC) curves for the CATH20 dataset and for various sequence redundancy levels of the SCOP-Lo dataset, along with their corresponding area under the curve (AUC) values. The all *vs*. all similarity calculation for the 3,673 proteins in the CATH20 dataset took 250 milliseconds. For the SCOP-Lo dataset, query *vs*. target similarity calculation for the 83 query proteins against the 10% target dataset (with 4,332 proteins) took 4 milliseconds and the 100% target set (with 23,912 proteins) took 20 milliseconds. Generating the target dataset embeddings was also fast, taking 2 minutes for the CATH20 dataset and 15 minutes for the 100% SCOP-Lo dataset (excluding file parsing time as this depends on the speed of the disk). Target set embedding time is not as important as search time, as it only has to be run once. Embedding each additional query protein takes 20-60 milliseconds depending on its length. A *k*-nearest neighbors classification of the CATH20 dataset into the five CAT classes showed a high accuracy of 82%.

**Fig. 3.**
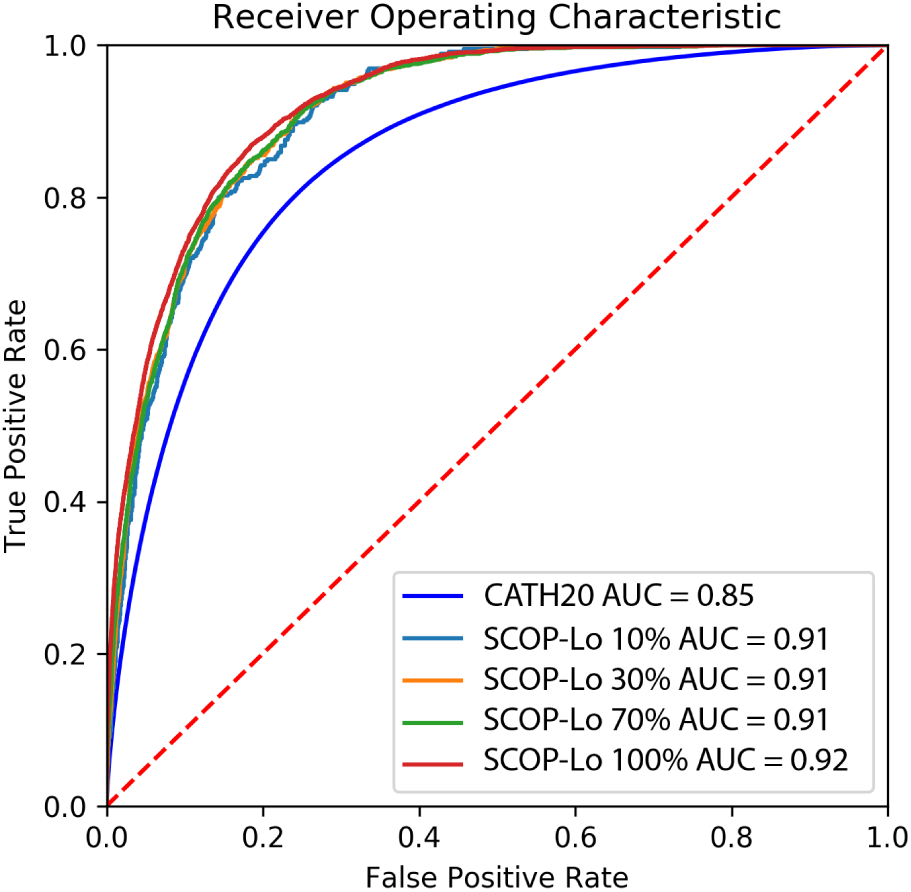
ROC curves for an all vs. all structure similarity search using Geometricus embeddings on the CATH20 dataset (dark blue) and four similarity searches of 83 query proteins on different sequence redundant target sets from the SCOP-Lo dataset (10% - light blue, 30% - orange, 70% - green, and 100% - red). True positives were determined using CATH and SCOP classifications, as described in the Methods.

In both these applications, Geometricus performs favourably compared to results reported by other alignment-free approaches applied to comparable datasets (Le *et al*., 2009) (Lo *et al*., 2007; Budowski-Tal *et al*., 2010), which typically achieve search AUCs between 0.75 and 0.85 and fold classification accuracy up to 75%. For the structural alphabets defined by Le *et al*. (2009) classification accuracy increases to 80% upon using more sophisticated SVM classifiers with tailored kernels. This approach is not investigated here but would likely improve our fold classification accuracy further. Geometricus comes close to the highly accurate alignment-based methods (Le *et al*., 2009) (with search AUCs exceeding 0.9 and fold classification accuracy exceeding 90%) at a mere fraction of the computational cost.

### 3.3 Geometricus can be used across and within protein families

Unlike library-based or deep learning-based structure embedding techniques, Geometricus can be adapted to the type and scale of the problem at hand without sacrificing speed, via the *m* (resolution) parameter. When comparing proteins from different superfamilies, a coarse discretization of structural fragments is preferred as it is expected that these proteins will have very different structures. However, as the specificity of the problem increases, the proteins under investigation start resembling each other more. In such cases, more specific binning of fragments, *i*.*e* a higher resolution, is advantageous to better capture their differences. This is demonstrated in Figure 4 with the Pfam10, CMGC, and MAPK datasets.

**Fig. 4.**
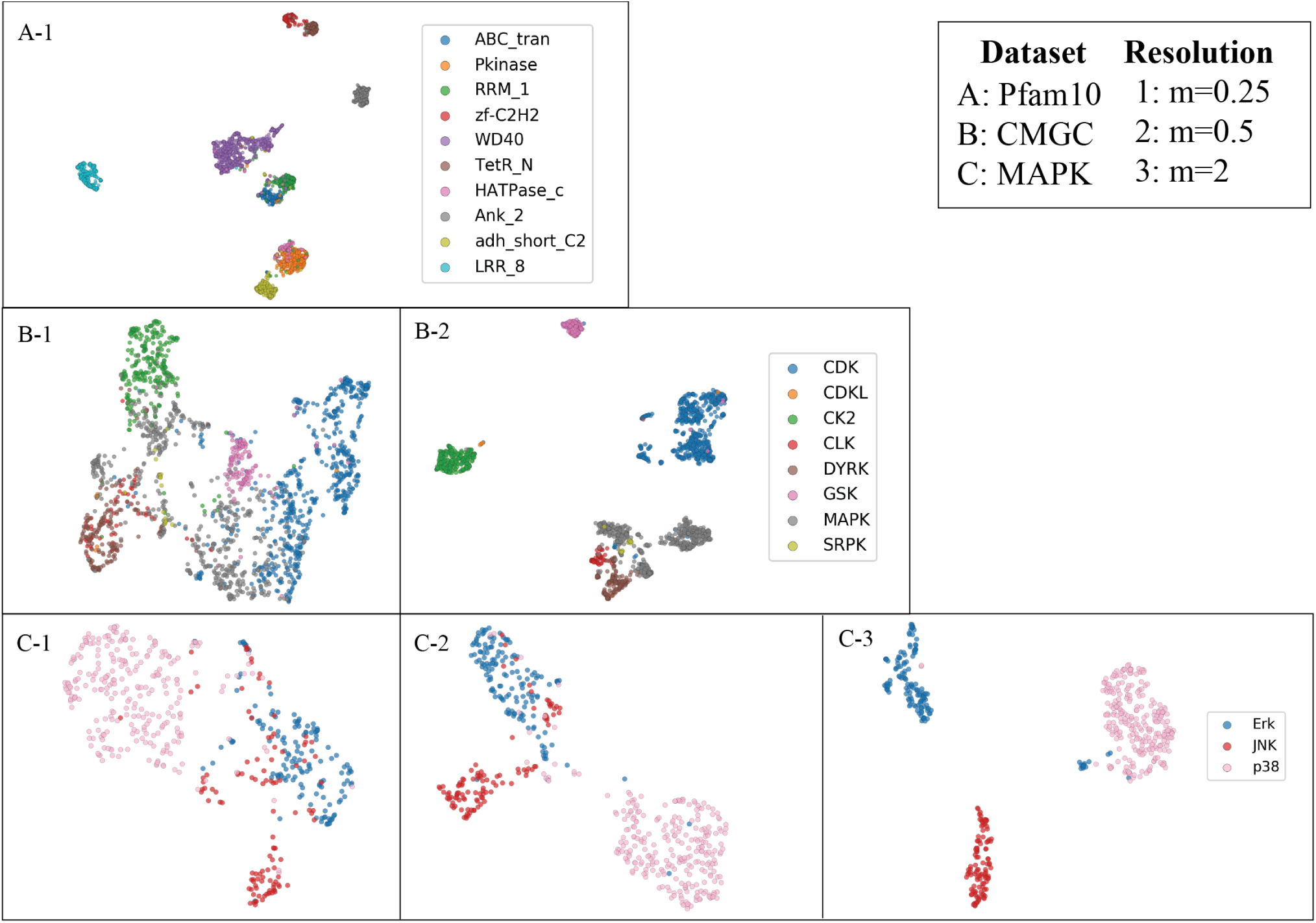
The effect of the resolution parameter *m* on different datasets. A, B, and C represent the Pfam10, CMGC, and MAPK datasets respectively. 1, 2, and 3 represent resolutions of 0.25, 0.5, and 2. As the structural similarity between proteins increases, higher resolutions are needed to achieve a good separation of pre-existing clusters in each dataset.

The Pfam10 dataset (Figure 4A) consists of proteins that contain one of ten fairly divergent Pfam domains. A low resolution of 0.25 (leading to 26 *k*-mer based shape-mers and 28 radius-based shape-mers) already separates these ten Pfam accessions into well-defined clusters. Some similar accessions, such as the Protein kinase domain (Pkinase) and the Histidine kinase-like ATPase (HATPase_c) cluster closer together as expected. Higher resolutions perform worse on this dataset, as comparable structural fragments are split across different shape-mers.

The CMGC dataset (Figure 4B) contains proteins from a group of kinases called the CMGC group (named after the initials of some members). As these proteins are more evolutionarily and functionally related, a higher resolution of 0.5 (resulting in 78 *k*-mer shape-mers and 93 radius shape-mers) is required to achieve a good separation between the individual families within this group.

Finally, the MAPK dataset (Figure 4C) consists of MAP kinases, a family of proteins which relay signals from the cell surface to coordinate growth, stress and other responses. This family is divided into subfamilies, here simplified into the p38, ERK, and JNK categories, each of which relay different types of growth and stress signals. A high resolution of 2 (resulting in 1098 *k*-mer shape-mers and 908 radius shape-mers) separates these subfamilies.

Thus, the feature space generated by Geometricus can be altered depending on the structural similarity expected between the proteins under consideration. This is especially advantageous in situations where the proteins under study are from the same family or subfamily and share a common structural fold, or in the case of mutation studies where local structure alterations occur due to single residue changes. In contrast, other embedding techniques are often optimized for divergent structures, and would likely assign the same embedding to each protein in these cases.

### 3.4 Geometricus can be used as input for interpretable machine learning

Typically, when analysing highly similar proteins as found in the MAPK dataset, one would also be interested in interpreting the results to find functionally important residues or structural regions. Such insights can be directly be applied to select candidate residues for mutational studies or used in directed evolution techniques to engineer proteins and enzymes with desired properties such as substrate specificity (Ding *et al*., 2014), drug-target binding affinity (Michael *et al*., 1992), interaction specificity (Fariselli *et al*., 2002), or thermostability (Jia *et al*., 2015) among others. Geometricus embeddings are well-suited for this kind of learning as each element of an embedding can easily be mapped back to the specific residues of the shape-mer it represents.

We demonstrate this with a classification problem defined for the MAPK dataset, namely to predict the specific subfamily of a MAP kinase. A simple decision tree trained on 70% of the data and tested on the remaining 30% showed an accuracy of 96% for this task. More interestingly, this trained classifier can now be inspected for predictive features. We mapped the top two shape-mers considered the most predictive by the decision tree back to all the residues and locations at which they occur across all the MAPK proteins. These locations are visualized on three example proteins, one from each of the three subfamilies (Figure 5A; shape-mer 1 in red, shape-mer 2 in blue). Figure 5B details the percentage of proteins from each subfamily which contain each of the two shape-mers, and the average number of times they appear per protein. The first appears more often in p38 kinases at a higher frequency per protein, while the second favors the ERK kinases with over three occurrences per protein on average. Looking at the structures themselves, it becomes clear which particular locations (highlighted and magnified) cause this difference in frequencies, even in such highly similar structures. While this is a simple example, it demonstrates the potential for using Geometricus in interpretable machine learning tasks for protein families.

**Fig. 5.**
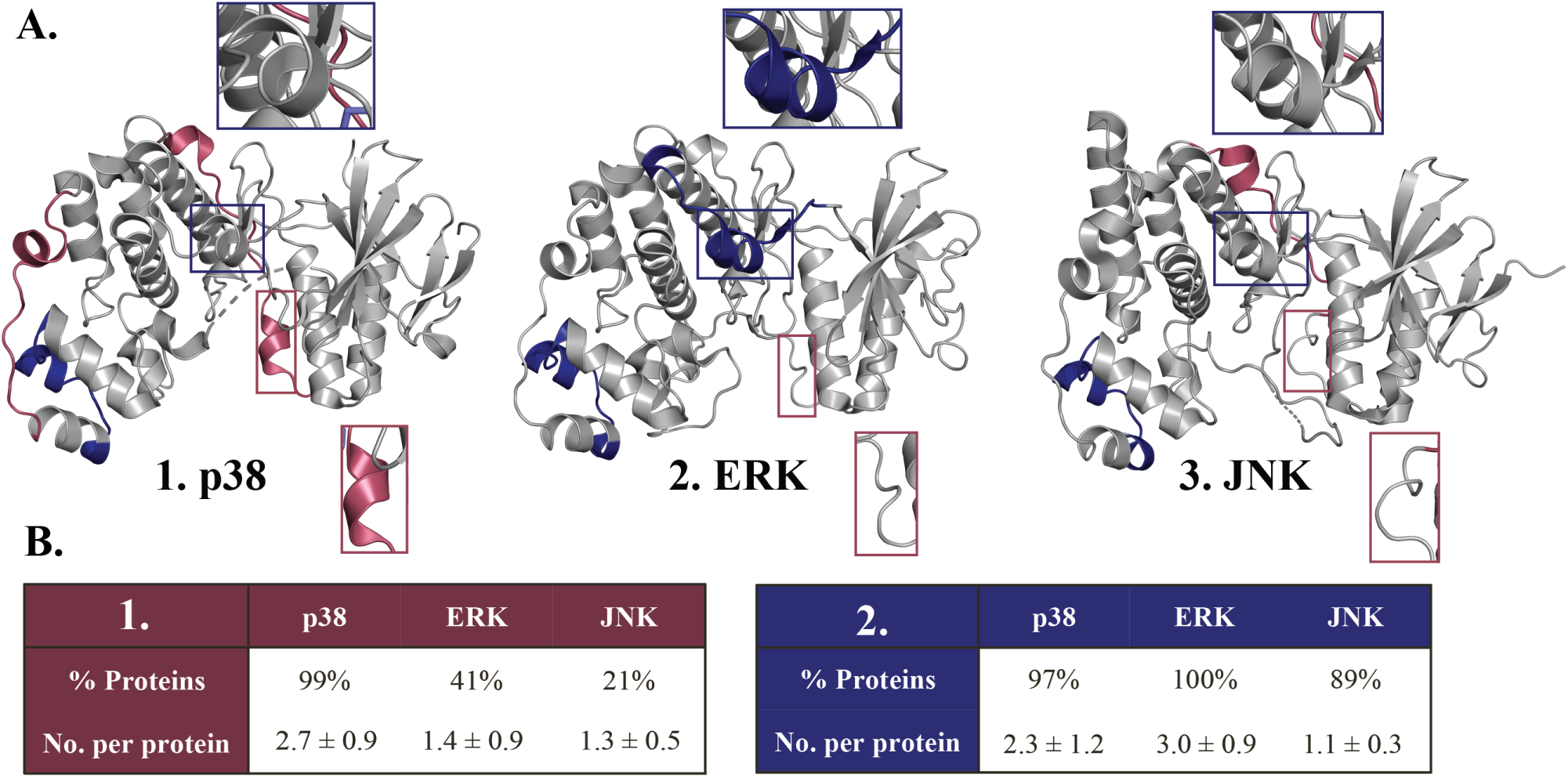
A. The occurrences of two shape-mers (colored red and blue respectively) most predictive in separating MAPK subfamilies visualized on three MAPK structures: 1. p38 structure (PDB ID: 3QUE), 2. ERK structure (PDB ID: 2OJJ) and 3. JNK structure (PDB ID: 4KKG). For each shape-mer, one location in a structure where it is present is magnified across all three structures and discussed in the text. B. The percentage of proteins containing a shape-mer and the average number of times a shape-mer appears per protein across the three MAPK subfamilies, for 1. the first shape-mer (red) and 2. the second shape-mer (blue).

## 4 Conclusion

We have presented a novel, fast and accurate approach for protein structure embedding with a wide range of applications. Geometricus uses 3D rotation invariant moments to describe structural fragments such that they can be easily compared across proteins without the need for superposition or alignment. This allows for a blazing fast embedding technique that takes milliseconds to generate an embedding of a protein, and scales linearly with the number of proteins.

The simplicity of this approach also brings with it versatility, as Geometricus does not depend on a fixed library of predefined fragments and can instead grow or shrink depending on the scale of the problem at hand. Therefore, it is readily applied to more specialised prediction tasks focusing on a single protein family with a conserved structural fold where other structure embedders would likely struggle to resolve each protein. The explicit mapping between residues and shape-mers further allows the user to trace back from a predictive model to predictive residues and structural regions, which can broaden our understanding of specific protein and enzyme mechanisms. This makes Geometricus well-suited for machine learning tasks where interpretation is a concern along with accuracy.

While this initial version of Geometricus uses four rotation-invariant moments, more such invariants have been studied (Žuni ć*et al*., 2016) and could be added to increase the specificity of a shape-mer. Another possible extension is to include solvent accessibility or amino acid descriptors as rotation-invariant aspects of a residue set. While these additions would likely not be so helpful in tasks spanning diverse proteins, such as structure similarity search, they may be useful in tasks involving enzyme mechanisms (Heckmann *et al*., 2018) or protein/ligand interactions and hotspots (Liu *et al*., 2018a; Zheng *et al*., 2019) where the accessibility of a structure fragment as well as its physicochemical and electrostatic properties matter as much as its shape.

Geometricus thus combines a set of highly attractive features that sets it apart from other structure embedding and structure similarity techniques. It is much faster than alignment-based algorithms such as Madej *et al*. (2014) and Ye and Godzik (2004), and at the same time highly accurate compared to other alignment-free techniques such as Le *et al*. (2009) and Lo *et al*. (2007). Unlike most techniques, its independence from a fragment library or predefined training set allows for broad application to generate feature sets for machine learning, even for differentiating mutants - something that has not been explored due to the focus of current techniques on divergent proteins. The shape-mer approach allows for easy interpretability and possible association of specific shapes to function, and its simplicity allows for ease of expansion. Shape-mer similarity could also be utilized to train structure-informed sequence embedding techniques, similar to the approach detailed by Bepler and Berger (2019), or as part of a scoring function to assess protein model quality, a field in which topology has been shown to play a crucial role (Garg *et al*., 2016).

Improvements in homology and *de novo* modelling techniques have greatly expanded the number of proteins for which we can accurately model structure. This means that future structure-based machine learning tasks will likely be augmented with structural models to obtain large datasets comparable to those used in sequence-based predictive approaches, where such a fast and versatile structural embedder would be useful. Given the prominent role in present-day bioinformatics of both structural modelling and machine learning, Geometricus embeddings, with possible further embellishments, may lead to breakthroughs in understanding protein function.

## Acknowledgements

We thank Rens Holmer and Barbara Terlouw for valuable feedback on drafts, stimulating discussions and insightful suggestions on the approach and applications presented here.

## References

Alley, E. C. et al. (2019). Unified rational protein engineering with sequence-based deep representation learning. Nature Methods, 16(12), 1315–1322.

AlQuraishi, M. (2019). ProteinNet: a standardized data set for machine learning of protein structure. BMC Bioinformatics, 20(1), 311.

Bakan, A. et al. (2011). Prody: protein dynamics inferred from theory and experiments. Bioinformatics, 27(11), 1575–1577.

Bateman, A. et al. (2002). The Pfam protein families database. Nucleic Acids Research, 30(1), 276–280.

Bepler, T. and Berger, B. (2019). Learning protein sequence embeddings using information from structure. arXiv preprint 1902.08661.

Bernstein, F. C. et al. (1977). The Protein Data Bank: A computer-based archival file for macromolecular structures. European Journal of Biochemistry, 80(2), 319–324.

Budowski-Tal, I. et al. (2010). FragBag, an accurate representation of protein structure, retrieves structural neighbors from the entire pdb quickly and accurately. Proceedings of the National Academy of Sciences, 107(8), 3481–3486.

DeLano, W. L. et al. (2002). PyMOL: An open-source molecular graphics tool. CCP4 Newsletter on Protein Crystallography, 40(1), 82–92.

Ding, H. et al. (2014). Similarity-based machine learning methods for predicting drug-target interactions: a brief review. Briefings in Bioinformatics, 15(5), 734–747.

Fariselli, P. et al. (2002). Prediction of protein–protein interaction sites in heterocomplexes with neural networks. European Journal of Biochemistry, 269(5), 1356–1361.

Flusser, J. and Suk, T. (1994). Affine moment invariants: a new tool for character recognition. Pattern Recognition Letters, 15(4), 433–436.

Flusser, J. et al. (2003). Moment forms invariant to rotation and blur in arbitrary number of dimensions. IEEE Transactions on Pattern Analysis and Machine Intelligence, 25(2), 234–246.

Garg, S. et al. (2016). Improved protein model ranking through topological assessment. In Computational Biology and Bioinformatics, pages 410–428. CRC Press.

Heckmann, D. et al. (2018). Machine learning applied to enzyme turnover numbers reveals protein structural correlates and improves metabolic models. Nature communications, 9(1), 1–10.

Hu, M.-K. (1962). Visual pattern recognition by moment invariants. IRE Transactions on Information Theory, 8(2), 179–187.

Jia, L. et al. (2015). Structure based thermostability prediction models for protein single point mutations with machine learning tools. PloS One, 10(9).

Kabsch, W. (1976). A solution for the best rotation to relate two sets of vectors. Acta Crystallographica Section A: Crystal Physics, Diffraction, Theoretical and General Crystallography, 32(5), 922–923.

Kooistra, A. J. et al. (2016). KLIFS: a structural kinase-ligand interaction database. Nucleic Acids Research, 44(D1), D365–D371.

Kratz, S. and Rohs, M. (2011). Protractor3D: a closed-form solution to rotation-invariant 3D gestures. In Proceedings of the 16th International Conference on Intelligent User Interfaces, pages 371–374.

Lam, S. K. et al. (2015). Numba: A LLVM-based python JIT compiler. In Proceedings of the Second Workshop on the LLVM Compiler Infrastructure in HPC, pages 1–6.

Le, Q. et al. (2009). Structural alphabets for protein structure classification: a comparison study. Journal of Molecular Biology, 387(2), 431–450.

Liu, S. et al. (2018a). Machine learning approaches for protein–protein interaction hot spot prediction: Progress and comparative assessment. Molecules, 23(10), 2535.

Liu, Y. et al. (2018b). Learning structural motif representations for efficient protein structure search. Bioinformatics, 34(17), i773–i780.

Lo, W.-C. et al. (2007). Protein structural similarity search by Ramachandran codes. BMC Bioinformatics, 8(1), 307.

Ma, J. and Wang, S. (2014). Algorithms, applications, and challenges of protein structure alignment. In Advances in Protein Chemistry and Structural Biology, volume 94, pages 121–175. Elsevier.

Madej, T. et al. (2014). Mmdb and vast+: tracking structural similarities between macromolecular complexes. Nucleic acids research, 42(D1), D297–D303.

Mamistvalov, A. G. (1998). N-dimensional moment invariants and conceptual mathematical theory of recognition n-dimensional solids. IEEE Transactions on Pattern Analysis and Machine Intelligence, 20(8), 819–831.

Mangin, J.-F. et al. (2004). Brain morphometry using 3D moment invariants. Medical Image Analysis, 8(3), 187–196.

McInnes, L. et al. (2018). UMAP: Uniform Manifold Approximation and Projection for Dimension Reduction. ArXiv e-prints.

Michael, J. et al. (1992). Modelling the structure and function of enzymes by machine learning. Faraday Discussions, 93, 269–280.

Moult, J. et al. (2016). Critical assessment of methods of protein structure prediction: Progress and new directions in round XI. Proteins: Structure, Function, and Bioinformatics, 84, 4–14.

Murzin, A. G. et al. (1995). SCOP: a structural classification of proteins database for the investigation of sequences and structures. Journal of Molecular Biology, 247(4), 536–540.

Pearl, F. M. et al. (2003). The CATH database: an extended protein family resource for structural and functional genomics. Nucleic Acids Research, 31(1), 452–455.

Pedregosa, F. et al. (2011). Scikit-learn: Machine learning in Python. Journal of Machine Learning Research, 12(Oct), 2825–2830.

Rao, R. et al. (2019). Evaluating protein transfer learning with TAPE. In Advances in Neural Information Processing Systems, pages 9686–9698.

Rizon, M. et al. (2006). Object detection using geometric invariant moment.

Sadjadi, F. A. and Hall, E. L. (1980). Three-dimensional moment invariants. IEEE Transactions on Pattern Analysis and Machine Intelligence, (2), 127–136.

Se, S. et al. (2001). Vision-based mobile robot localization and mapping using scale-invariant features. In Proceedings 2001 ICRA. IEEE International Conference on Robotics and Automation (Cat. No. 01CH37164), volume 2, pages 2051–2058. IEEE.

Senior, A. W. et al. (2020). Improved protein structure prediction using potentials from deep learning. Nature, pages 1–5.

Simossis, V. et al. (2003). An overview of multiple sequence alignment. Current Protocols in Bioinformatics, 3(1), 3–7.

Sommer, I. et al. (2007). Moment invariants as shape recognition technique for comparing protein binding sites. Bioinformatics, 23(23), 3139–3146.

Ye, Y. and Godzik, A. (2004). Fatcat: a web server for flexible structure comparison and structure similarity searching. Nucleic acids research, 32(suppl_2), W582–W585.

Zheng, N. et al. (2019). Targeting virus-host protein interactions: Feature extraction and machine learning approaches. Current drug metabolism, 20(3), 177–184.

Žunić, J. et al. (2016). On a 3D analogue of the first Hu moment invariant and a family of shape ellipsoidness measures. Machine Vision and Applications, 27(1), 129–144.

